# Mechanical loading of the Achilles tendon during different rehabilitation exercises: A cross sectional observational study

**DOI:** 10.1101/2023.06.13.544659

**Authors:** Paul New, Lianne Wood

## Abstract

**Background:** Achilles tendon injuries are common in active populations and heel raising exercises are commonly used in tendon rehabilitation programmes. This study compared characteristics of force occurring within the healthy Achilles tendon during two different types of exercise conditions that can be used in Achilles tendon rehabilitation.

**Method:** A cross-sectional, observational study was conducted to compare force fluctuations occurring within the Achilles tendon during two different types of heel raising exercise. All subjects performed firstly, a set of traditional eccentric heel drops (HD) and secondly an adapted walking drill (WD), and results were compared. 13 Healthy subjects were recruited from staff and post graduate students as a sample of convenience from the biomechanics department at the University of Bath. Tendon forces were calculated using a combination of data collected from force plate, motion analysis and real time ultrasound.

**Results:** Fluctuations in force were seen in all subjects in both exercise conditions. The HD condition produced a statistically significant increase in force fluctuations compared to the WD condition (p<0.001).

**Conclusion:** This study shows that force fluctuations can be stimulated at different levels during functional movement patterns and exercise conditions. Therefore, exercise routines can be tailored to meet individual needs and the stage of the pathology. These findings may have implications for exercise progressions in the rehabilitation of Achilles tendon disorders. Future research is needed determine if differences in tendon force fluctuations correlate with pathology.

**Summary Statement:** This study is of interest to clinicians, exercise professionals and researchers with an interest in the study of Achilles tendon biomechanics and the management of tendon injuries.

## Introduction

Achilles Tendinopathy (AT) is a common debilitating injury among both athletic and non-athletic populations, with an annual adult incidence rate of 2.35 per 1000 (Malvankar and Khan, 2011). 36% of those who suffer an episode never return to their previous level of activity (de Jonge, et al., 2011; Miller et al., 2021). The condition is also a common finding in metabolic disorders such as obesity, diabetes and hypercholesterolaemia (Ackerman and Renstrom, 2012). 44% of those who undertake treatment are left with ongoing pain and dysfunction at long term follow up (Miller et al., 2021).

The disorder is notoriously unresponsive to clinical intervention (Arya and Kulig, 2010). The aetiology is poorly understood and research explaining how exposure to mechanical loads result in cellular changes in tendons tissue is limited (Magnusson and Kjaer, 2019). However, it is widely accepted that a combination of anatomical and biomechanical factors combine in the presence of repeated excessive mechanical strain to cause micro-trauma to the tendon (Van Usen and Pumberger, 2007; Miller et al, 2021).

### Anatomy

The Achilles tendon is made up of the combined insertions of the Gastrocnemius and Soleus muscles (Fig. 1) to form the Triceps Surae (Norkin and Levangie, 1983). Through its attachment to the calcaneal bone, the tendon facilitates the transfer of muscular force to control movement of the body over the foot (Wyndow et al., 2010; Reinking, 2012). The elastic recoil properties of the tendon produce a spring-like action during locomotion which increases efficiency of movement (Litchwark and Wilson, 2005).

**Fig. 1.**
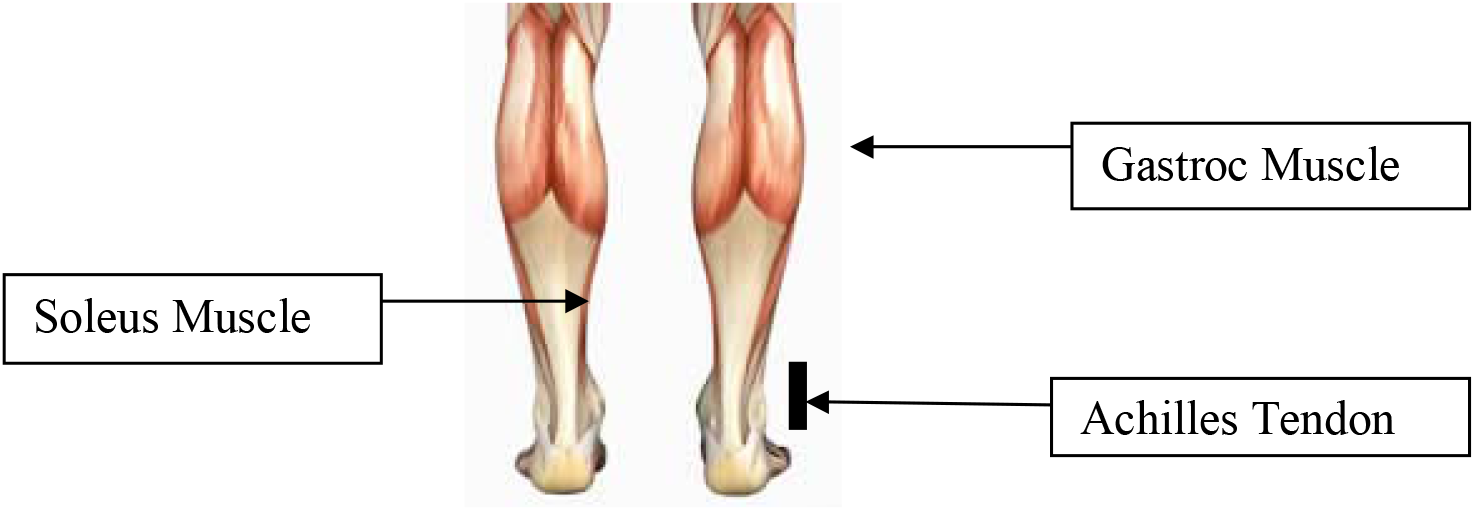
The Anatomy of the Triceps Surae.

The tendon consists predominantly of type I and type III collagen fibres that are arranged in a tightly packed parallel formation. The fibres give a dense hierarchical protein structure (Fig. 2), with elastic properties, that are suitable for tolerating the high tensile forces of fast locomotion (Malvankar and Khan, 2011; Warden and Scott, 2011). Healthy tendon tissue consists of spindle shaped tendon cells (tenocytes) interspersed with extra cellular matrix (ECM) (Benjamin et al., 2008). Tenocytes are responsible for the secretion of the ECM and the turnover of collagen to maintain tendon structural properties (Cook et al., 2002; Benjamin et al., 2008: Riley, 2008; Parkinson et al., 2011).

**Fig. 2.**
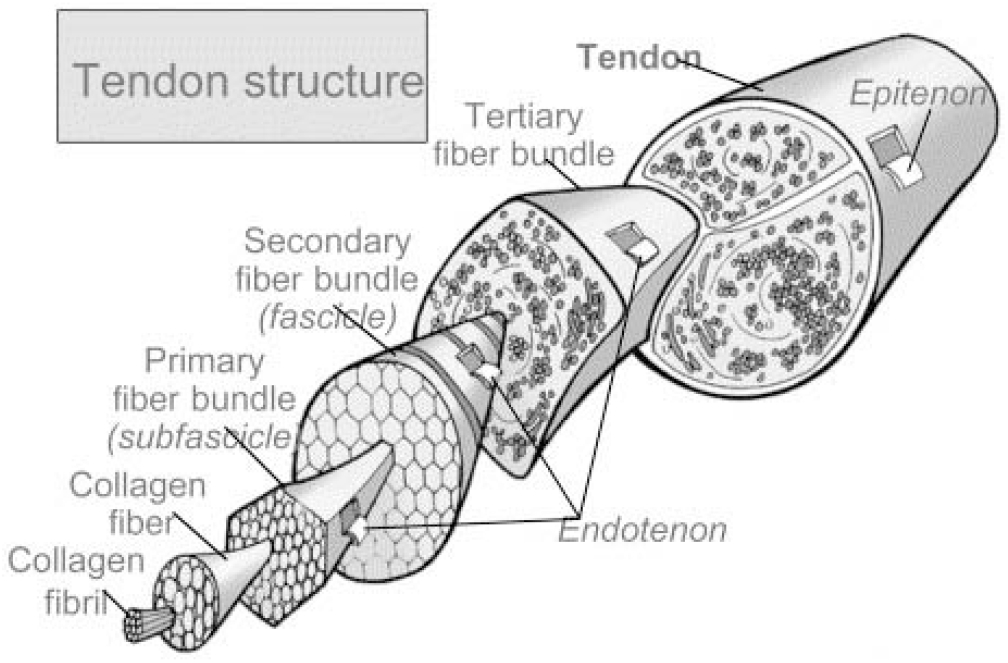
Tendon hierarchial structure Ashe. M. C. & McCauley, T. (2004). Tendinopathies in the upper extremity: A paradigm shift. *Journal of Hand Therapy*, 17, 3, 329–334.

### Pathology

Tendinopathy implies a load induced pathological change within the tendon tissue (Wyndow, Cowan, Wrigley, & Crossley, 2010). Rees, Maffulli and Cook (2009), propose that changes occur within the material structure in a clinical syndrome characterised by pain, swelling, and impaired function. Histologically the injured tendon displays disorganisation of collagen fibres and hypercellularity within the ECM (Ayra and Kulig, 2010; Fung et al., 2010; Ackerman and Renstrom, 2012), resulting in a weaker, less organised structure with a lower failure level (Po-Yee et al., 2009; Nevasier et al., 2012). As the condition progresses, tissue changes produce pain and swelling in what is considered to be a failed healing or degenerative response within the tendon (Wyndow et al., 2010).

### Treatment

Interventions should focus on the reversal of tendinopathic changes and not just the resolution of symptoms (Miller et al., (2021). Animal models have demonstrated that low levels of intermittent exercise can influence tendon adaptation by increasing anabolic growth factors to promote a healing response (Eliasson et al., 2012; Eliakim et al., 2000). Remodelling is stimulated through a process of ‘mechano-transduction’ where forces are transformed into cellular responses, via a complex series of biological signalling pathways (Kahn and Scott, 2009; Martino et al., 2018; Wang and Linlin, 2022). At a macro level, optimal mechanical loading induces changes in material properties, including stiffness and load tolerance (Kjaer et al., 2009; Kingma et al., 2007; Childs et al, 2010; Heinemeier and Kjaer, 2011; Silbernagel et al., 2020; Wang and Linlin, 2022).

Stanish, Rubinovich and Curwin (1986) first introduced the rationale that eccentric muscle contractions were an effective way to stimulate a therapeutic response. This approach was adapted specifically for AT in a land mark study by Alfredson, Pietilla and Jonsson (1998), who demonstrated a decrease in pain and a return to activity using eccentric heel drop exercises. However, the rationale led to the presumption that isolated eccentric forces were required to stimulate the repair process (Miller et al., 2021. This was refuted by later research where all types of muscle contraction were shown to be of benefit (Beyer et al., 2015; Littlewood et al., 2015). The assumption that high forces are needed to remodel the tendon may be inaccurate and all types of muscle contraction are now thought to be useful in rehabilitation (Rees et al., 2008; Morrisey et al., 2011; Malliaras et al., 2015). Other authors have reported that eccentric loading regimes maybe painful, resulting in the pursuit of other treatment options (Maffulli et al., 2008; Silbernagel et al., 2020). The material properties of the injured tendon have been shown to be biologically and mechanically inferior at 6 weeks post injury with tendon microstructure often not fully restored to pre-injury status (Freedman et al., 2018). It is recognised that tendons heal through the formation of scar tissue, with increased cartilage deposits, altered tenocyte shape and raised cell numbers, reflecting a tendon with a reduced capacity to transfer strain (Zhang et al., 2016; Freedman et al., 2018).

Optimal loading conditions for tendon remodelling remain unknown and current trends in clinical research literature suggest that interventions should be individually tailored to meet personal needs (Silbernagel et al., 2020). In healthy subjects, studies have shown that load can be progressively increased by advancing through incremental exercise levels, (Rowe et al., 2012) and it has become widely accepted that pain should be used as a guide for succession (Silbernagel et al., 2007; Silbernagel et al., 2020). Conversely, when assessed in a longitudinal cohort study of 24 male runners with mid portion AT, pain responses where highly variable and did not corelated with load progression in 73% of the cohort (Sancho et al., 2023). However, this small study was not a randomised controlled trial, making it difficult to extrapolate findings to infer cause and effect but it does highlight the unexplained relationship that pain plays in tendinopathy. Smith et al., (2017) propose that discomfort may not be a barrier to recovery as moderate quality evidence exists to support the benefits of working into pain during exercise protocols for chronic musculoskeletal conditions. This poses potential problems for clinical practice by questioning the reliability of pain guided exercise progressions in the rehabilitation of AT.

Clinical improvements observed with therapeutic loading routines may be due to sinusoidal force fluctuations occurring within the tendon, rather than the magnitude of the force produced (Rees et al., 2009). Fluctuations in muscle force result from the challenge of controlling a lower limb dynamic movement (Litchwark and Wilson, 2005). AT is characterised by a reduced capacity within the tendon to tolerate load (Grigg et al., 2014). Lower force frequencies have been observed in a small single cohort study of all male participants with AT, when compared to controls (Grigg et al., 2013). These force frequency fluctuations seen in tendon tissue, have been proposed as a stimulus for cellular remodelling through mechano-transduction (Rees, et al., 2008; Seynnes et al., 2009). Increased force fluctuations, which occur in faster or more challenging movements, result from the difficulty of recruiting large motor units in the muscle tissue. These oscillations stimulate adaptive cellular responses within the tenocytes themselves and influence structural and material changes in tendon properties (Chaudry and Screen, 2011).

Research supports the progression from low to high load exercises, with the aim of promoting tissue tolerance to mechanical strain (Wang and Linlin, 2022). Sancho et al., (2023) identified that commonly prescribed AT rehabilitation exercises, as described by Silbernagel, Hanlon and Sprague (2020) fall into 2 distinct clusters defined by low and high load intensity. The current approach promoted by clinical guidelines (ICON, 2019) implies the assumption that a linear relationship exists between tendon pain and increasing load. However, Freedman et al., (2018) inform that tendon responses to load are not linear and interestingly found that lower loads resulted in more collagen fibre realignment in injured tendons.

It seems plausible that tendinopathic tissue may need longer periods, working at lower loads to enable sufficient adaptive remodelling to occur (Arampatzis et al., 2007: Silbernagel et al., 2007; Rowe et al., 2012). The aim of this study was to examine force fluctuations in two different weightbearing Achilles tendon specific exercises to determine if the tendon can be challenged at different levels. This may allow for a more gradual transition between low and high load exercise progressions in tendon rehabilitation programmes.

## Method

Characteristics of force fluctuations within the Achilles tendon were compared in two different exercise conditions. A walking drill (WD) exercise was adapted from a weight bearing neuromuscular control protocol described by Elphinston (2008) to challenge the tendon micro-environment and was compared with results during a traditional eccentric heel drop routine (HD) (Alfredson et al., 1998). It was hypothesised that similar inversions of the force trace could be stimulated in the WD as that produced in a HD.

### Participants

13 healthy volunteers (mean ± sd age = 35 ± 4 years; height = 1.76 ± 0.09m; mass 76 ± 11.5 kg; male 9 (69.2%) and female 4 (30.8%) were recruited into the study as a sample of convenience in a similar cohort to that described by Rees *et al*., (2008). We recruited participants from staff and post graduate students attending the Biomechanics laboratory at the University of Bath during the summer semester. A word of mouth recruitment strategy was used to enrol subjects, who were free from lower limb musculoskeletal injury with no previous history of AT. Written informed consent was obtained from each subject and eligibility granted if aged over 18 and free from any other medical or lower limb disorder. Ethical approval was granted by the School for Health Research Ethics Committee at the University of Bath and complied with the regulations set out in the Declaration of Helsinki (2008).

### Data collection

A cross-sectional observation study was conducted to track force fluctuations within the Achilles tendon using inverse dynamics analysis. This is a mathematical method of calculating internal forces and joint moments from the measurements of limb motion, tendon moment arm and ground reaction forces to provide information on the tendon’s mechanical work. (Farris et al., 2008) This allows an estimation of Achilles tendon force during movement (Rees et al., 2008). All data was collected in the biomechanics laboratory at the University of Bath.

The instantaneous moment arm of the Achilles tendon was calculated via the position of the tendon vector, from the muscle-tendon junction (MTJ), to the calcaneus and the centre of rotation of the ankle joint (a virtual mid-point between the two malleoli) by utilising a combination of CODA motion analysis (Charnwood Dynamics, UK) and real time ultrasound in B mode (UAB Telemed, Lithuania) (Litchwark & Wilson, 2005). The ultrasound probe was taped to the lower limb over the medial Gastrocnemius, at the myotendinous junction (MTJ), using elasticated adhesive bandage to image the position of the MTJ at 50Hz. A CODA motion analysis system recorded the position of an active marker set placed on the ultrasound probe and anatomical landmarks on the lower limb, including the calcaneus, 5^th^ MTP joint, lateral and medial malleoli and head of fibula (Farris et al., 2011; 2012). 3D Marker coordinates were calibrated in an embedded laboratory reference frame as described by Litchwark and Wilson (2005).

The exercises where performed on a force plate (Kisler Instruments Ltd, Switzerland) and ground reaction data was recorded at 1000Hz. This method has been used to good effect in previous tendon research and has been shown to be reliable in assessing the mechanical properties of the tendon in functional movements (Litchwark and Wilson, 2005; Rees et al., 2008; Farris, 2009). The technique has been used to examine force fluctuations within the Achilles tendon during eccentric exercise by Rees et al., (2008). Estimations of tendon force were made from the data every 0.02 seconds in the load bearing phase of the exercises (McGuigan et al., 2007).

Force fluctuations were deemed as inversions of the force trace, with two inversions or more per exercise cycle considered significant, as has been defined in previous research (Rees et al., 2008). An inversion was recorded when the force value dropped and then climbed again during the loading phase of each exercise cycle.

### Exercise protocol

Subjects were asked to perform one set of 15 repetitions of two different eccentric exercises (HD and WD) in a random order. Each subject was given instruction by a trained physiotherapist (lead author) on how to perform both exercises and given 5 minutes to practise both conditions (HD and WD). Participants then performed a 1 x 15 repetition set of each exercise condition and data was acquired from the last five repetitions of each condition, as in previous research (Rees et al., 2008). Each repetition was performed to a count of one second up (concentric phase) and three seconds down (eccentric phase) for all 15 repetitions in both exercise conditions. Subjects were allowed a 5 minutes rest period in between the two trials. The peak Achilles tendon force and the number of inversions in the force-time trace were calculated in each repetition for 1 set of both exercise conditions.

All data was analysed using MATLAB™ (The Mathworks Inc, USA) computer software and descriptive statistics were calculated for each exercise across the group of participants. Results were compared for within group and between groups (HD vs WD) differences. Statistical significance was analysed using a paired t-test and the level of statistical significance set at P = 0.005.

## Results

Sinusoidal fluctuations in Achilles tendon loading were observed in both exercise conditions for all subjects. An increasing rise in Achilles tendon force was also seen in the eccentric phase of both conditions as subjects lowered their heel from full plantar-flexion to end of range dorsiflexion. Higher peak force values and a greater angle of ankle dorsi-flexion were observed in the HD group (Fig. 5).

**Fig. 3.**
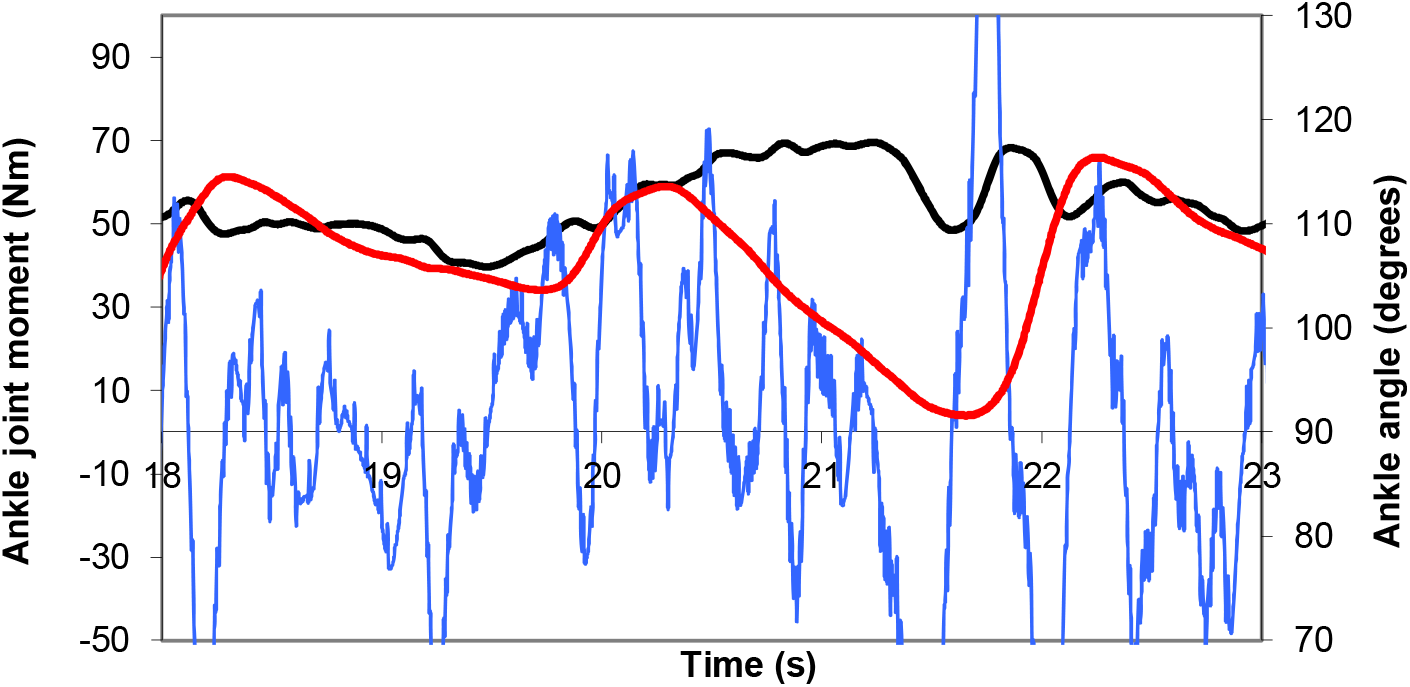
A. Force fluctuations during an adapted Walking Drill (WD) exercise. Black line = Tendon force trace. Redline = Ankle joint angle. Blue line = Fluctuations in force

**Fig. 3.**
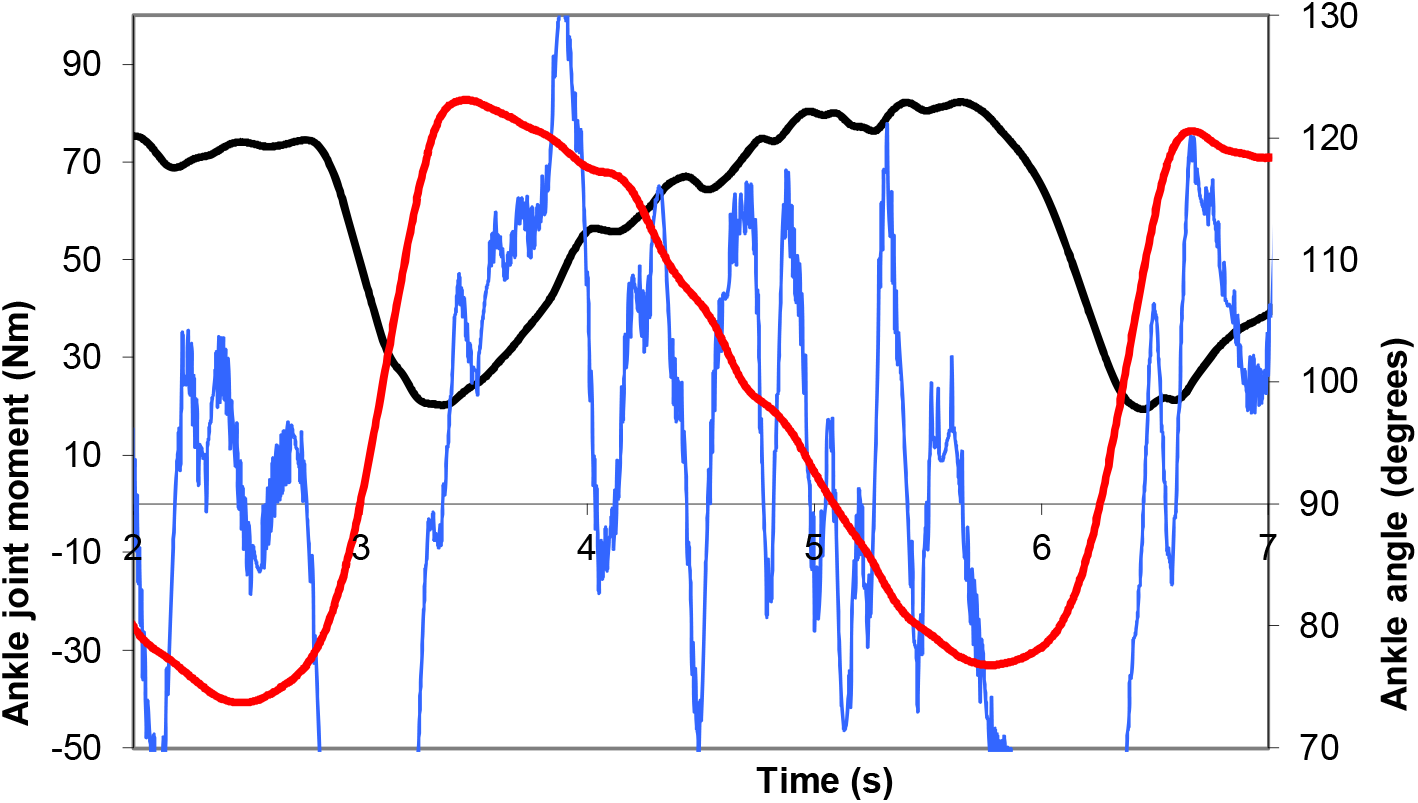
B. Force Fluctuations during a Heel Drop (HD) exercise. Black line = Tendon force trace. Redline = Ankle joint angle. Blue line = Fluctuations in force

**Fig. 4.**
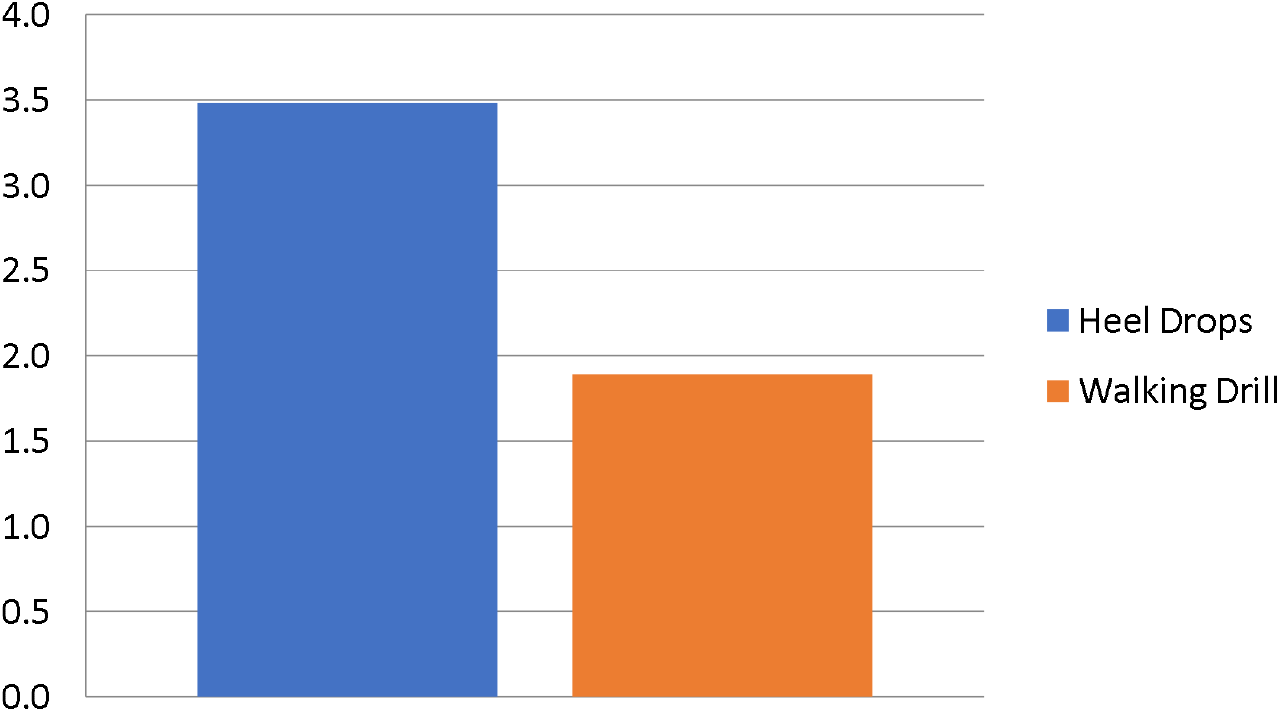
Graph to show mean difference in force fluctuation between the two exercise conditions.

**Fig. 5.**
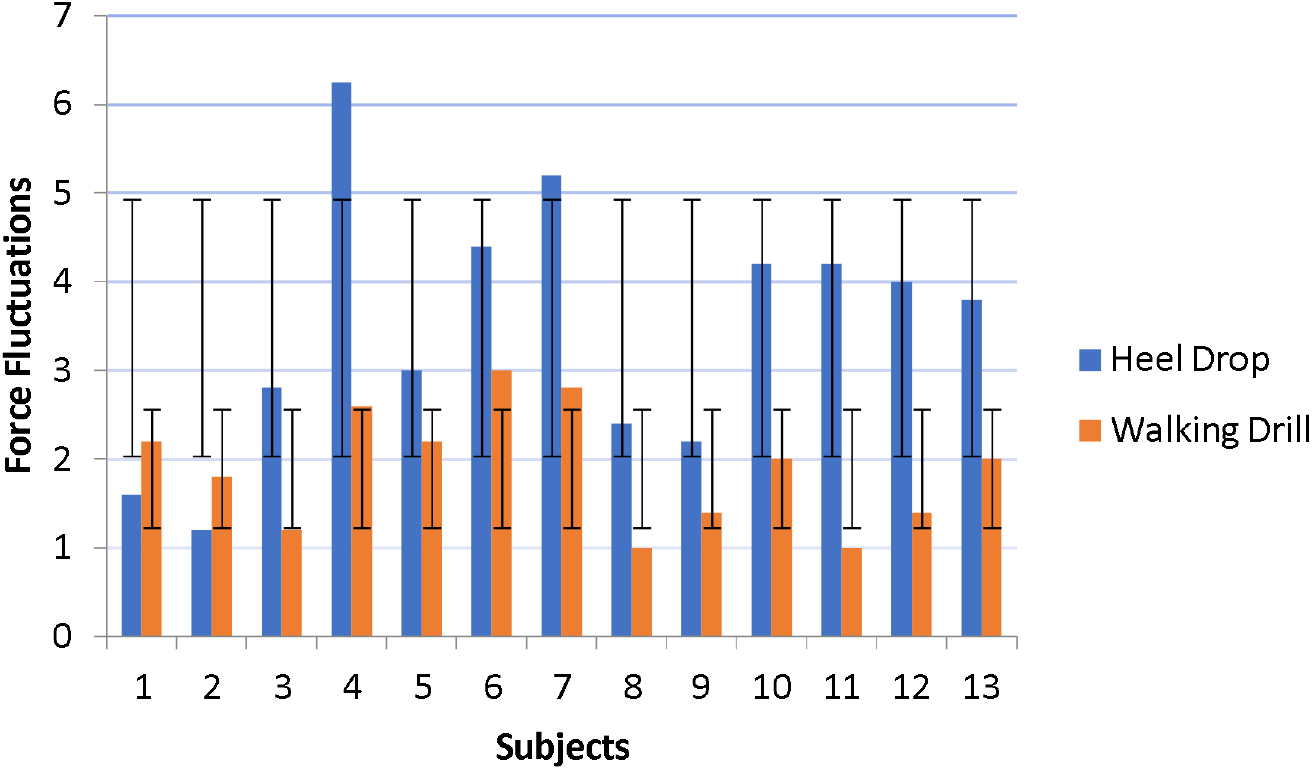
Graph to show individual force fluctuations in all subjects during both exercise conditions.

All subjects displayed frequent fluctuations of the force trace during the eccentric phase in both conditions. On average two (mean 1.9, SD 0.7) force fluctuations per cycle were recorded in the WD, while four fluctuations (Mean 3.5, SD 1.9) per cycle were seen in the HD exercise (Fig. 6).

The difference between the two conditions was statistically significant P = 0.0007. Individual force fluctuations are shown in Figure 7. Analysis shows that a higher individual variance in force fluctuation was produced in the HD condition, which is reflected in the greater standard deviation seen in this exercise condition.

## Discussion

This study investigated differences in force fluctuations occurring in the healthy Achilles tendon during two different exercise routines (HD and WD). Results showed that force fluctuations arise during the eccentric phase of both exercise conditions. The main finding was that subjects displayed a statistically significant increase in force fluctuation in the HD condition compared to the WD. These force fluctuations have been proposed as a mechanism for the effectiveness of eccentric loading exercises in the rehabilitation of AT (Rees et al., 2008). Our results support the hypothesis that variability in force characteristics within the healthy Achilles tendon can be stimulated at different levels in different exercise conditions.

In keeping with previous research, findings show that greater force fluctuations are stimulated by the eccentric phase of an exercise, confirming such oscillations are a characteristic of eccentric movements (Rees et al., 2008; Morrissey et al., 2012; Henrikson et al., 2011). Rees et al., (2008) speculate that this mechanism possibly results from the difficulty of controlling a dynamic movement. Henrikson et al., (2009) found similar results using EMG and suggest that a vibration above 10 Hz may cause small changes in neural activation which stimulate fibroblasts to form increased gap junctions and cross links between tendon fibres. An increase in neuro-peptides has been reported after eccentric exercise, leading to an increase in blood flow and the normalisation of tendon structure (Scott and Bahr, 2009). This indicates that the remodelling process may well be stimulated by mechanical force fluctuations rather than the force magnitude or the type of muscle contraction.

In this study, episodes of loading and unloading within the Achilles tendon occurred at twice the frequency during the HD condition (Mean = 4 per cycle) compared to the WD (Mean = 2 per cycle). This indicates that the HD posed a greater challenge to neuromuscular control and therefore represents a greater mechanical stimulus to the Achilles tendon. The HD condition offered a smaller base of support and required subjects to perform an eccentric movement to a greater range of motion than the WD, which may explain this finding.

These results suggest that the WD exercise may be an appropriate intervention to include earlier in a rehabilitation plan, where adaptations need to be stimulated initially, at a lower level without causing pain (Kulig et al., 2008; Reiman and Lorenz, 2011; Rutland et al., 2010). These results support the work of Silbernagel et al., (2020) who propose that AT rehabilitation plans should be progressive in load application. Knowledge of force fluctuations in different exercises would allow a smoother transition between the cluster groups of exercises identified by Sancho et al., (2023). While loading does need to be of an optimal type and intensity to encourage a therapeutic response, exercise progression in rehabilitation remains problematic (Anderson et al., 2009; Davenport and Kulig., et al., 2005; Fung et al., 2010; Rutland et al., 2010). Silbernagel et al., (2020) advocate that pain should be used as a marker for advancement, however the work of Sancho et al., (2023) has cast doubt on the notion that pain and load are corelated in a linear relationship. Therefore, while empirical evidence suggests that mechanical reloading should occur in a step wise manner, it may not be the case that pain can be used as a marker for progression without accelerating tendon pathology.

Contemporary AT rehabilitation programmes focus on stimulating changes in tendon structure and ECM adaptations without consideration of the sensorimotor or behavioural aspects of human movement. The current aim of rehabilitation is to reduce tendon compliance by enhancing mechanical properties without causing further damage (Childs et al., 2010). Theoretical models of muscle/tendon physiology are largely based on unproven assumptions that force fluctuations or variability arise out of noise in the process of converting central nervous system drive into muscular force, producing kinematic variability (Nagamori et al., 2021). Such random variations in force output increase in amplitude with the level of input and the challenge presented in controlling a dynamic movement. These variations also reduce with repeated exposure to the stimulating level of physical activity. Nagamori et al., (2021) suggest that this may represent more than just noise within the nervous system.

The limitations of this study are that it was only conducted in a small group of subjects free from tendon pathology, that were recruited from a single centre as a convenience sample. Therefore, these results are not generalisable to a wider healthy population or those with AT. No repeated measures analysis was conducted, limiting the reliability of results. The study methodology utilised inverse dynamics, as opposed to direct tendon measurements, and therefore does not evaluate the contributing actions of other muscle groups, such as Tibialis Anterior. This may lead to an over estimation of load placed on the Achilles tendon (Rees et al., 2008; Litchwark et al., 2005; Farris, 2009).

Future research should examine force fluctuations in the AT population to determine if characteristics correlate with pathology, providing useful information to guide rehabilitation progressions. It is currently unknown how force variability, produced in dynamic movements effects the remodelling of tendon pathology. More research into therapeutic exercise is needed to determine if force fluctuations can be modified over time to enhance tendon mechanical properties and guide rehabilitation in AT patients. Research conducted in animal models has shown that individuals respond to load differently and a threshold would appear to exist where optimal mechanical stress stimulates anabolic effects within the ECM (Arampatzis et al., 2007; Eliasson et al., 2012). This study suggests that, exercise routines should be tailored to meet individual needs while considering the stage of the pathology and the demands of the required activity (Andarawis-Puri et al., 2012). If the reliability of pain as a marker for progression in rehabilitation is uncertain (Sancho et al., 2023), then it seems reasonable to hypothesise that force fluctuations may play a more important role in tendon loading than previously thought (Nagamori et al., 2021).

## Conclusion

This study compared force characteristics in the Achilles tendon in two different exercise routines. The main findings show that HD exercises load the Achilles tendon to a statistically significant greater level than WD’s. Therefore, force fluctuations can be gained during functional movement patterns and stimulated at different levels within the Achilles tendon, implying that tendon loading routines can be tailored to individual needs, as recommended by current guidelines. Future research should examine if these findings are repeatable in a population with tendon pathology.

## Author Contribution

Concept, idea, research design & writing: P. New Consultation and review: L. Wood

## Acknowledgements

This study was completed as part of an MSc in Sports Physiotherapy that was conducted at the University of Bath in 2013. The lead author would like to thank the following: Dr Polly McGuigan for her expert advice and excellent supervision which made completing this project possible and great fun. Dr Lianne Wood for lending her time, helpful advice and clear guidance on the writing up of this paper. Portsmouth Football Club for its financial support and commitment to advancing professional development in elite sport.

